# Single-cell profiling coupled with lineage analysis reveals distinct sacral neural crest contributions to the developing enteric nervous system

**DOI:** 10.1101/2022.05.09.491197

**Authors:** Weiyi Tang, Jessica Jacobs-Li, Can Li, Marianne E. Bronner

## Abstract

During development, the enteric nervous system (ENS) arises from neural crest cells that emerge from the neural tube, migrate to and along the gut, and colonize the entire intestinal tract. While much of the ENS arises from vagal neural crest cells that originate from the caudal hindbrain, there is a second contribution from the sacral neural crest that migrates from the caudal end of the spinal cord to populate the post-umbilical gut. By coupling single cell transcriptomics with axial-level specific lineage tracing in avian embryos, we compared the contributions between embryonic vagal and sacral neural crest cells to the ENS. The results show that the two neural crest populations form partially overlapping but also complementary subsets of neurons and glia in distinct ganglionic units. In particular, the sacral neural crest cells appear to be the major source of adrenergic/dopaminergic and serotonergic neurons, melanocytes and Schwann cells in the post-umbilical gut. In addition to neurons and glia, the results also reveal sacral neural crest contributions to connective tissue and mesenchymal cells of the gut. These findings highlight the specific properties of the sacral neural crest population in the hindgut and have potential implications for understanding development of the complex nervous system in the hindgut environment that may influence congenital neuropathies.

---

The enteric nervous system (ENS), the largest component of the peripheral nervous system, plays a critical role in regulating gut motility, hormone homeostasis, as well as interactions with the immune system and gut microbiota^1^. In amniotes, the ENS consists of millions of neurons with motor, sensory, secretory and signal transduction functions as well as a larger number of enteric glia. Together they form a highly orchestrated network of physically and chemically connected cells embedded between the muscle and mucosal layers of the gastrointestinal system^2^. Due to the ENS’s vast number of neurons and glia and its capacity for autonomic regulation, it is often referred to as “a second brain”^3^.

The neurons and glia of the ENS originate from the neural crest, a migratory stem cell population that forms most of the peripheral nervous system. During development, these cells migrate from the central nervous system (CNS) into the periphery to colonize the intestines. Much of the ENS is derived from “vagal” neural crest cells that arise in the caudal hindbrain, enter the foregut and migrate from caudally to populate the entire length of the gut. The vagal neural crest arises in the CNS adjacent to somites 1-7 at Hamburger Hamilton (HH) stage 10 and fully colonizes the entire length of the gut by embryonic day (E) 8 in the chick embryo^4^, after undergoing one of the longest migrations of any embryonic cell type. However, there is an additional neural crest contribution to the ENS from a neural crest population that arises at the caudal end of the CNS, referred to as the “sacral” neural crest. First observed by Le Douarin and Teillet in quail-chick chimeric grafts, sacral neural crest cells enter the hindgut and migrate rostrally to contribute neurons and glia to the post-umbilical gut^4^. The sacral neural crest arises posteriorly to somite 28 at HH stage 17-18, fully colonizes the post-umbilical gut by E8 and expands in number by E10^5, 6^ (Fig 1A).

**Figure 1.**
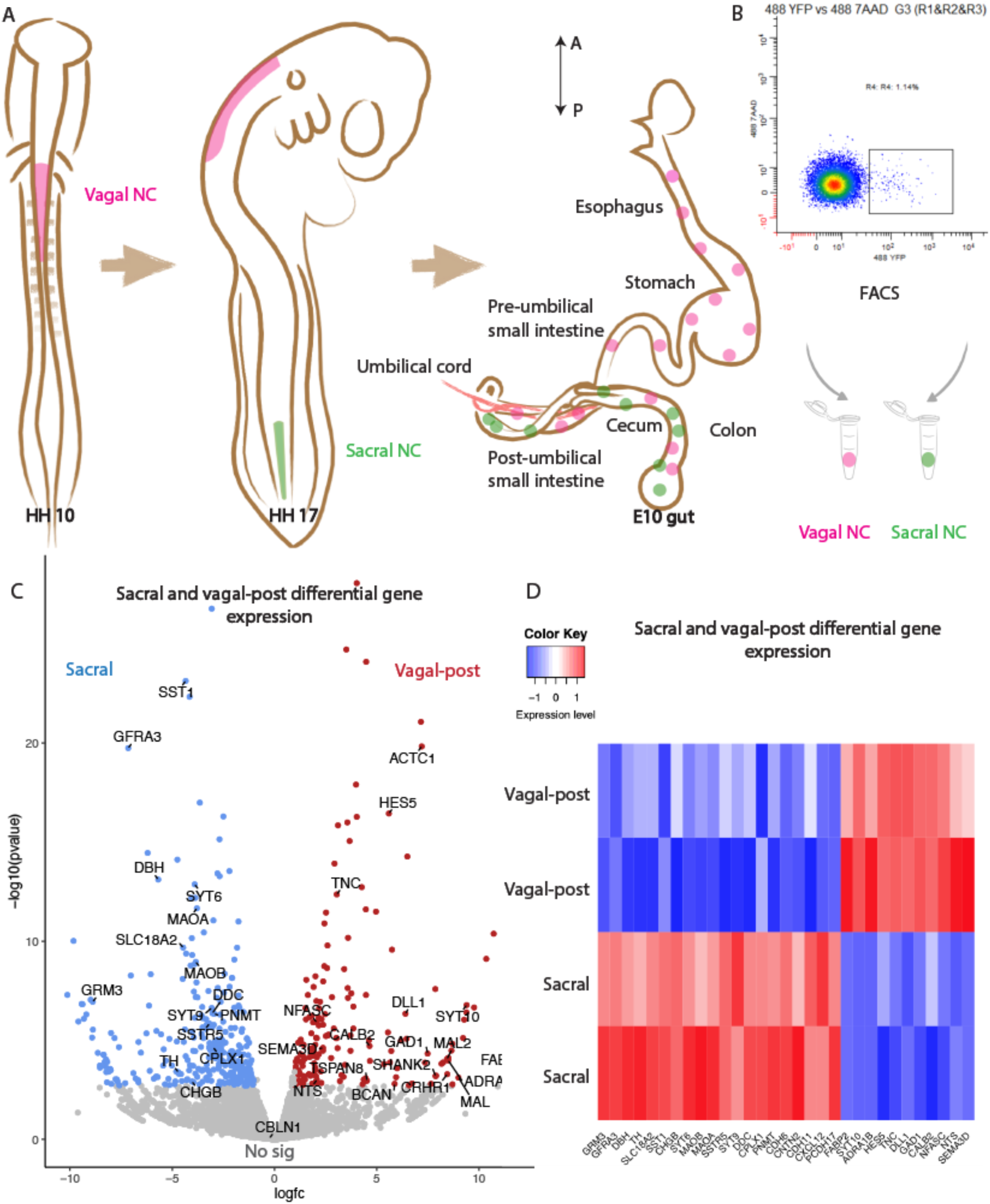
Comparative transcriptomic analysis of vagal and sacral neural crest derived cells in post-umbilical ENS. **(A)** Schematic diagram describing experimental procedure. Vagal (magenta) and sacral (green) neural crest cells were labelled by H2B-YFP in separate embryos. The post-umbilical gastrointestinal tracts were dissected at E10 for dissociation. From anterior (A) to posterior (P), the chicken gastrointestinal tract can be divided into six regions including esophagus (eso), stomach (sto), pre-umbilical small intestine (pre.int), post-umbilical small intestine (post.int), cecum (ce) and colon (col). **(B)** YFP+ cells from the post-umbilical region derived from vagal and sacral neural crest (NC) were sorted via FACS. **(C)** Volcano plot describing differentially expressed genes of sacral (blue) and vagal neural crest cells in the post-umbilical gut (vagal-post, red). Genes with fold change greater than 2 and p value less than 0.05 are colored. **(D)** Heatmap highlighting selected genes related to neuronal functions from differential gene expression analysis in sacral and vagal-post ENS populations (with 2 replicates per condition). Genes are ordered based on significance level and fold change.

Dysregulation of ENS development is responsible for enteric neuropathies including Hirschsprung’s disease which affects 1 in 5000 live births and is characterized by hindgut aganglionosis that results in the lack of gut motility^7^. While the etiology of Hirschsprung’s disease is not completely understood, it is thought that insufficient migration or proliferation of vagal neural crest precursors results in neuronal deficits, particularly in the hindgut due to the long distance needed for precursor cells to reach their destination. Grafting sacral neural crest cells in place of ablated vagal neural crest resulted in isolated ganglia in myenteric and submucosal plexuses. However, the grafted sacral neural crest’s contribution to the ENS was insufficient in number to compensate for the lack of vagal-derived cells, suggesting that there are intrinsic differences between these two populations^8^. Molecular cues such as GDNF^9^ and ET3^10^ are essential for migration and differentiation of vagal neural crest during early development. Consistent with this, mutations in *Ret*, *Gdnf*, *Ednrb* and *Edn3* genes are common in patients with Hirschsprung’s disease^11^, although phenotypes of these mutations often exhibit a complex inheritance pattern and low penetrance^7^.

Recent studies have proposed that lack of rostrocaudal migration of the vagal neural crest may not be the only cause of enteric neuropathies. A distinction between the ENS of the foregut/midgut versus hindgut is that the former arises solely from vagal neural crest-derived cells whereas the latter is populated by both vagal and sacral neural crest cells^5^. Thus, deciphering the ontogeny of hindgut neuropathy requires a thorough understanding of all neural crest-derived contributions to the hindgut and exploring possible cell fate diversity between vagal and sacral neural crest populations. Open questions include: what is the complete complement of cell types derived from the sacral neural crest? Is the sacral neural crest population distinct from the vagal, or do they have shared derivatives? Does the post-umbilical gut possess special neuronal cell types absent in the pre-umbilical region? A more complete view of the transcriptional landscape of the sacral compared with the vagal neural crest will provide a deeper understanding of these different cell populations and clarify how dysregulation of their developmental programs may contribute to congenital birth defects.

To tackle these questions, we combined single-cell transcriptomics with a recently developed lineage tracing method utilizing replication-incompetent avian (RIA) retroviruses to specifically label distinct neural crest cells populations in developing chick embryos. As amniotes, avian embryos closely resemble human embryos at similar developmental stages, while being much more accessible to experimental manipulation than mammalian embryos. RIA retroviral infection permanently expresses a fluorescence signal in the infected cell lineage^12^ without the need for transplantation or Cre-mediated recombination that can result in ectopic expression. This allows for precise region-specific lineage analysis to follow cell fate and isolate pure populations of cells derived from vagal or sacral population, thereby enabling transcriptional profiling at single cell resolution. Thus, the combined features of the chick embryo and the RIA retrovirus provide a particularly advantageous system to comparatively study the relative contributions of the vagal and sacral neural crest cells to the entire developing ENS for the first time.

By selectively labeling each neural crest population, we demonstrate that sacral neural crest cells form functionally distinct units from the vagal neural crest, suggesting that these populations are likely to have distinct functions. Single-cell transcriptome analysis shows that the sacral neural crest gives rise to the majority of Schwann cells and adrenergic/dopaminergic/serotonergic neurons of the post-umbilical gut, whereas the pre-umbilical vagal neural crest is the primary source of secretomotor neurons. Interestingly, vagal neural crest cells in the post-umbilical gut are intermediate in character between the pre-umbilical vagal and sacral populations, suggesting a strong influence of environmental cues in cell fate decisions. In addition, our results uncover sacral neural crest contributions to connective tissue and melanocytes in the post-umbilical gut region. This study expands our understanding of the sacral neural crest, a largely understudied stem cell population that acts in close coordination with the vagal neural crest to form essential neuronal and glial cells of the hindgut ENS. Our results further suggest that specific properties of vagal and sacral neural crest could be essential in understanding the nature of congenital hindgut neuropathies.

## Results

### Population level RNA-seq analysis shows that the sacral and vagal neural crest populations exhibit distinct transcriptional profiles

As a first step in assessing contributions of vagal and sacral neural crest in the post-umbilical region, we obtained pure populations of each using a novel replication-incompetent avian (RIA) retrovirus lineage tracing technique^12, 13^ for bulk RNA-seq analysis. To this end, the neural tube at the level of the caudal hindbrain was injected with RIA retrovirus carrying a YFP expression cassette at Hamburger Hamilton stage 10 (HH10) to label vagal neural crest cells, or below the level of somite 28 at HH17 to label the sacral neural crest. The post-umbilical guts were then harvested at embryonic day 10 (E10, Fig 1A) by which time the gut was fully populated by neural crest cells that were undergoing terminal differentiation; after dissociation, YFP+ cells were sorted using FACS (Fig 1B). Similar regions from three guts were pooled as a replicate, with each library containing 2000 cells.

Differential gene expression analysis reveals intriguing distinctions between vagal and sacral neural crest cells in the post-umbilical gut at the population level, suggesting that they serve distinct functions. Genes enriched in the sacral population include *Sst1/Sstr* indicating interneuron cell fate, and *Dbh, Th, Ddc, Pnmt,* and *Slc18a2* which are present in catecholaminergic neurons and serotonergic neurons. *Grm3* expression indicates that glutamatergic character is also more abundant in the sacral population. In addition, we observed up-regulation of *Gfra3* which is involved in the GDNF signaling pathway and *Cxcl12* which is related to signaling during cell migration (Fig 1C, D). Conversely, the vagal post-umbilical population expresses the adrenergic receptor *Adra1b*, enzyme *Gad1*, *Calb2* and *Nts* consistent with excitatory motor neuron fate. Additionally, the population expresses genes related to neural crest and neuronal migration such as *Sema3d* and *Tnc*. *Hes5 and Dll1* expression indicate functions for Notch/Delta signaling in the vagal-post-umbilical population (Fig 1C, D). These results suggest that vagal and sacral neural crest cells give rise to diverse cell types likely reflecting different functions^14, 15^.

### Single-cell transcriptome profiling of the chick ENS

To understand the transcriptional profile of vagal and sacral neural crest-derived cells at single cell resolution, we performed viral labeling as described above (Fig 1A) and collected three distinct neural crest populations at E10: vagal neural crest from the pre-umbilical gut (vagal-pre; 3 guts pooled per replicate), vagal neural crest from the post-umbilical gut (vagal-post; 6 guts pooled per replicate) and sacral neural crest from the post-umbilical region (sacral; 6 guts pooled per replicate). After FACS isolation of YFP+ cells, 4.6k-5k cells were sequenced for each replicate, generating a single-cell profile with ∼29000 cells that organized into 13 clusters^16–18^. To ascribe cluster identity, we first performed gene expression heatmap analysis for top 10 gene markers in clusters 0-12 (C0-C12), marking the most up-regulated genes as compared to all other clusters (Fig 2A).

**Figure 2.**
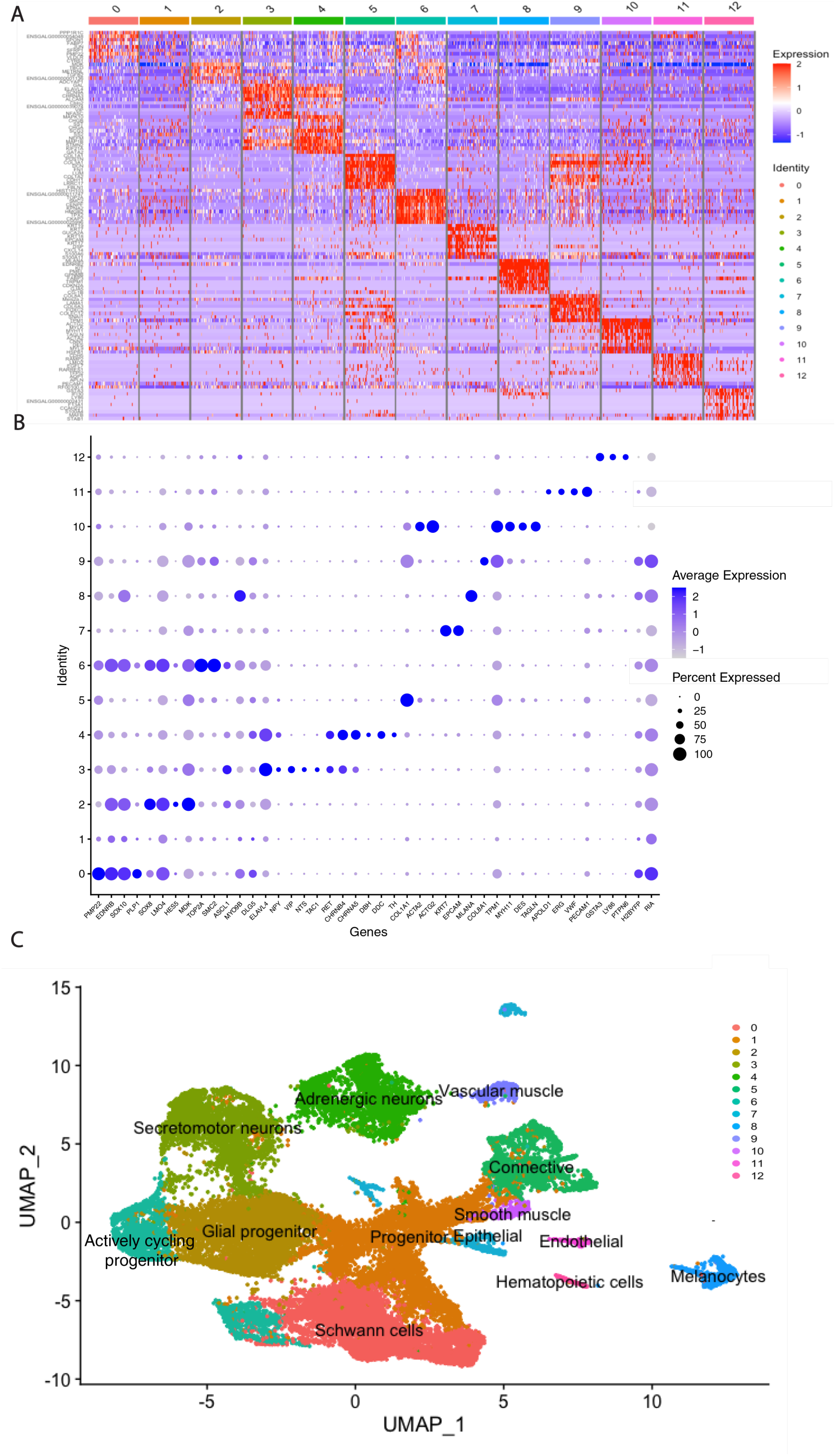
Single-cell transcriptomics of the post-umbilical ENS. **(A)** Expression heatmap for top 10 gene markers in clusters 0-12(subsampled). **(B)** Dotplot showing expression level and percentage of cell type specific markers (*Pmp22*, *Ednrb*, *Sox10*, *Plp1*, *Sox8*, *Lmo4*, *Hes5* glial progenitor marker; *Mdk*, *Top2a, Smc2* dividing cell marker; *Ascl1*, *Myo9b, Dlg5* neuronal precursors; *Elavl4*, differentiated neurons; *Npy*, inhibitory motor neurons; *Vip*, *Nts*, *Tac1*, excitatory motor neurons; *Ret*, enteric neuron marker; *Chrnb4*, *Chrna5*, cholinergic; *Dbh*, *<u>Ddc</u>*, *Th*, adrenergic; *Col1a1*, connective tissue; *Krt7*, *Epcam*, epithelial; *Mlana*, melanocytes; *Col8a1*, *Tmp1*, vascular muscle; *Acta2*, *Actg2*, *Myh11*, *Des*, *Tagln*, smooth muscle; *Apold1*, *Erg*, *Vwr*, *Pecam1*, endothelial; *Gsta3*, *Ly86*, *Ptpn6*, hematopoietic-related markers). Lineage tracer (H2B-YFP) and viral genome sequences (RIA) were examined across the clusters. Due to low YFP and RIA transcripts, C10 and C12 will not be a focus for downstream analysis. **(C)** Uniform manifold approximation and projection (UMAP) representation with cell identities assigned according to marker gene expression most represented in the cluster.

Next, we selected a group of genes characteristic of cell fate to define the dominate cell type in the clusters and present them as a dotplot (Fig 2B) and feature expression (Fig 3A). We identified C0 as Schwann cells based on high levels of *Pmp22*, *Ednrb*, *Sox10*, and *Plp1* expression (Fig 2B, Fig 3A *Sox10*, *Pmp22*, *Endrb*). Consistent with heatmap features, C1 is likely to contain progenitor cells due to its expression of *Myo9D*, which is associated with p75 signaling, and low expression of differentiation markers (Fig 2B). C2 exhibits high levels of the neural crest marker *Ednrb*, progenitor/glial marker *Sox10*, proliferation marker *Mdk*, as well as high levels of *Sox8*, *Lmo4* and *Hes5.* Due to this gene expression profile alongside the lack of Schwann cell markers, C2 was considered a glial progenitor population (Fig 2B). C3 and C4 have high expression of *Elavl4*, a marker for differentiated neurons, *Ret* representing enteric neurons, and the neuropeptide *Npy*, suggesting that both clusters have *Npy*+ inhibitory motor neurons (Fig 2B, Fig 3A *Ret*, *Elavl4*, *Npy*). However, C3 exhibits more prominent *Vip*, *Nts*, *Tac1* expression than C4, indicating excitatory motor neurons; C4 has more cholinergic genes like *Chrna5*, and adrenergic/serotonergic genes such as *Dbh*, *Ddc*, and *Th* (Fig 2B, Fig 3A). Therefore, C3 is likely to represent a secretomotor neuron cluster, while C4 is characteristic of adrenergic neurons, although neurons with other functions are also present. Undifferentiated and proliferating cells are most abundant in C2 and C6, as they exhibit the highest *Mdk* expression. Some cells in C2 and C6 express *Ascl1*, indicating they may be neuroblasts and/or glioblasts (Fig 2B, Fig 3A *Mdk, Ascl1*). However, C2 has glial markers like *Sox8* indicating it is a glial progenitor cluster. Due to high expression of cell cycle associated genes like *Top2a* and *Smc2*, progenitor cells in C6 seem to be actively cycling (Fig 2B).

**Figure 3.**
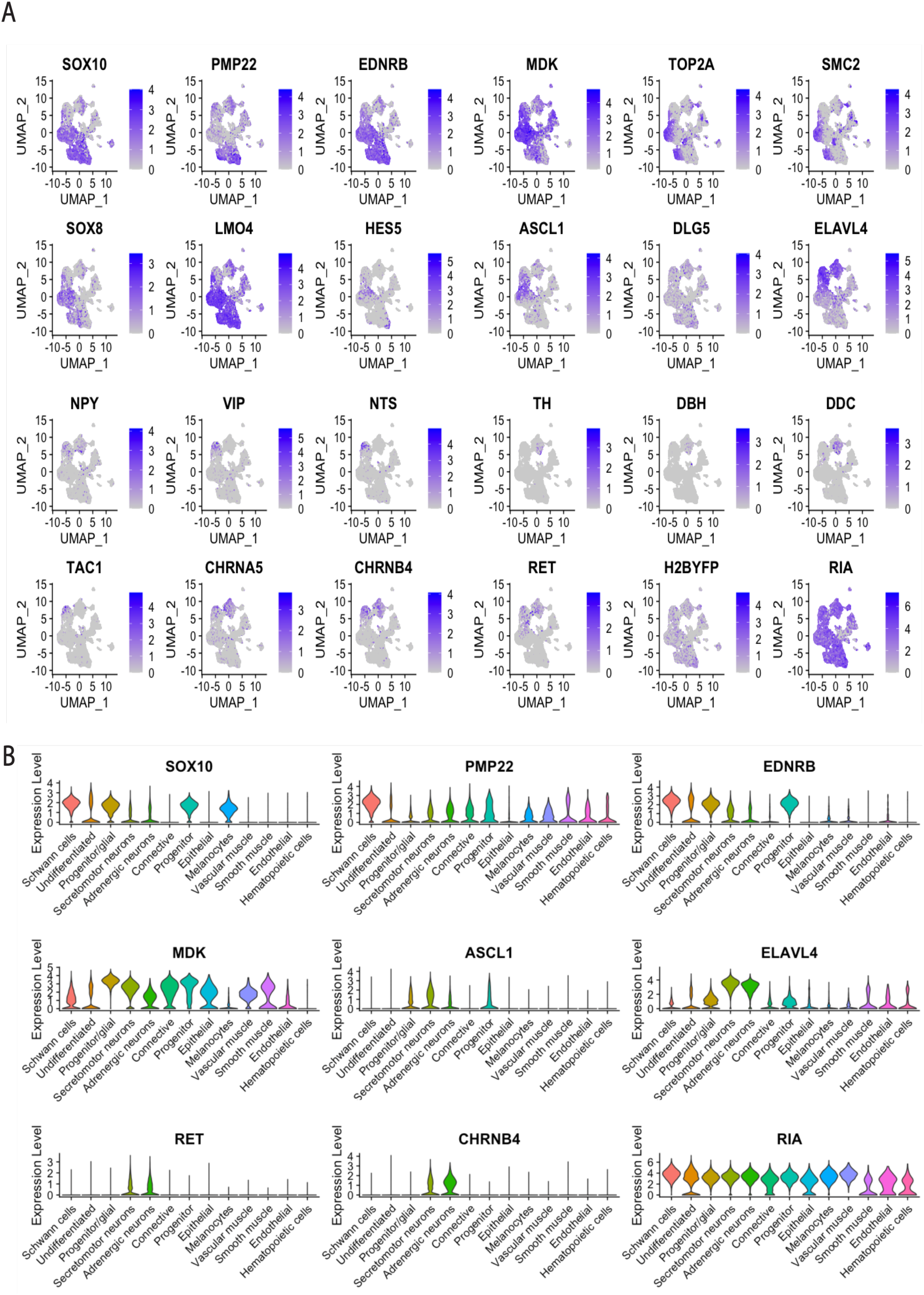
Gene expression analysis for ENS cell fates. **(A)** Feature plot of genes related to neuronal/glial/progenitor cell fates, neuronal transmitters and neuropeptides associated with specific neuronal subtypes. **(B)** Violin plot of key gene markers representing neuronal/glial/progenitor cell fates. *Ascl1*-expressing neuronal precursors are distributed in clusters related to neuronal cell fates, while ENS markers *Ret* and *Chrnb4* are restricted in differentiated neurons.

In addition to canonical peripheral nervous system derivatives, our analysis has revealed a range of cell fates previously unknown to be derived from the neural crest in the gut. For example, C5 is classified as connective tissue cells based on *Col1a1*. C7 is epithelial cells as suggested by markers *Krt7* and *Epcam*. Based on expression of *Mlana,* C8 is likely to be composed of melanocytes. C9 has a similar expression pattern to C5, but its *Col18a1* expression likely reflects a vascular muscle fate. C10 appears to be smooth muscle cells based on strong expression of *Acta2*, *Actg2*, *Tpm1*, *Myh11*, *Des*, and *Tagln*. C11 expresses endothelial markers *Apold1*, *Erg*, *Vwf*, and *Pecam*. C12 is likely to be hematopoietic-related cells based on *Gsta3*, *Ly86*, and *Ptpn6* expression (Fig 2B).

Putative cluster identities are presented in Uniform Manifold Approximation and Projection (UMAP, Fig 2C). To exclude the possibility that these cell types are a result of contamination during cell sorting, we examined lineage tracer (H2B-YFP) and viral genome sequences (RIA) expression across the clusters (Fig 2B, Fig 3A H2B-YFP, RIA, Fig 3B RIA). RIA genomic transcripts were abundantly expressed in almost all clusters, suggesting these were indeed virally-labeled neural crest derived cells. Some cells in the progenitor cluster (C1) exhibit lower YFP and RIA transcripts; these cells also showed lower expression for other markers (*Sox10*, *Ednrb*) except for moderate level of *Mdk*, possibly due to low overall transcription level. Because C10 and C12 generally lacked YFP and RIA transcripts, the two clusters are likely to be a result of autofluorescence, and thus were eliminated from downstream analysis.

### Vagal and sacral neural crest differentially contribute to ENS cell types

Next, we explored the relative contributions of the sacral, and pre- and post-umbilical vagal cell populations to this broad range of cell types present in the E10 gut. According to the UMAP separated by populations, sacral, post-umbilical vagal and pre-umbilical vagal populations form distinct combinations of cell types (Fig 4A, B). Quantification of cluster contributions reveals a common contribution to progenitor cells (C1), connective tissue (C5), actively cycling progenitors (C6), and epithelial cells (C7). In contrast, the remaining clusters have primary contribution from one or two populations (Fig 4C). For example, the Schwann cell cluster (C0) is mainly comprised of sacral (orange) and vagal (navy) populations in the post-umbilical gut with minimal contribution from the pre-umbilical vagal neural crest cells (light blue) (Fig 4C). This is likely to be a result of non-myelinating Schwann cells located in the intestinal nerve of Remak, present in the post-umbilical gut. The glial progenitor (C2) population is primarily derived from pre-umbilical vagal population. Interestingly, over 75% of C3 secretomotor neurons are from the vagal neural crest derived pre-umbilical gut, with a small proportion (20%) from the post-umbilical vagal population. However, adrenergic neurons in C4 were restricted to the post-umbilical gut. Most post-umbilical neurons are contributed to by the sacral neural crest. C8 melanocytes are only found in the post-umbilical region, with most cells contributed from sacral neural crest. C9 is a vascular smooth muscle population that almost exclusively arises from sacral neural crest. While endothelial cells are present throughout the entire gut length, with sacral and post-umbilical vagal populations as the major contributors (Fig 4C).

**Figure 4.**
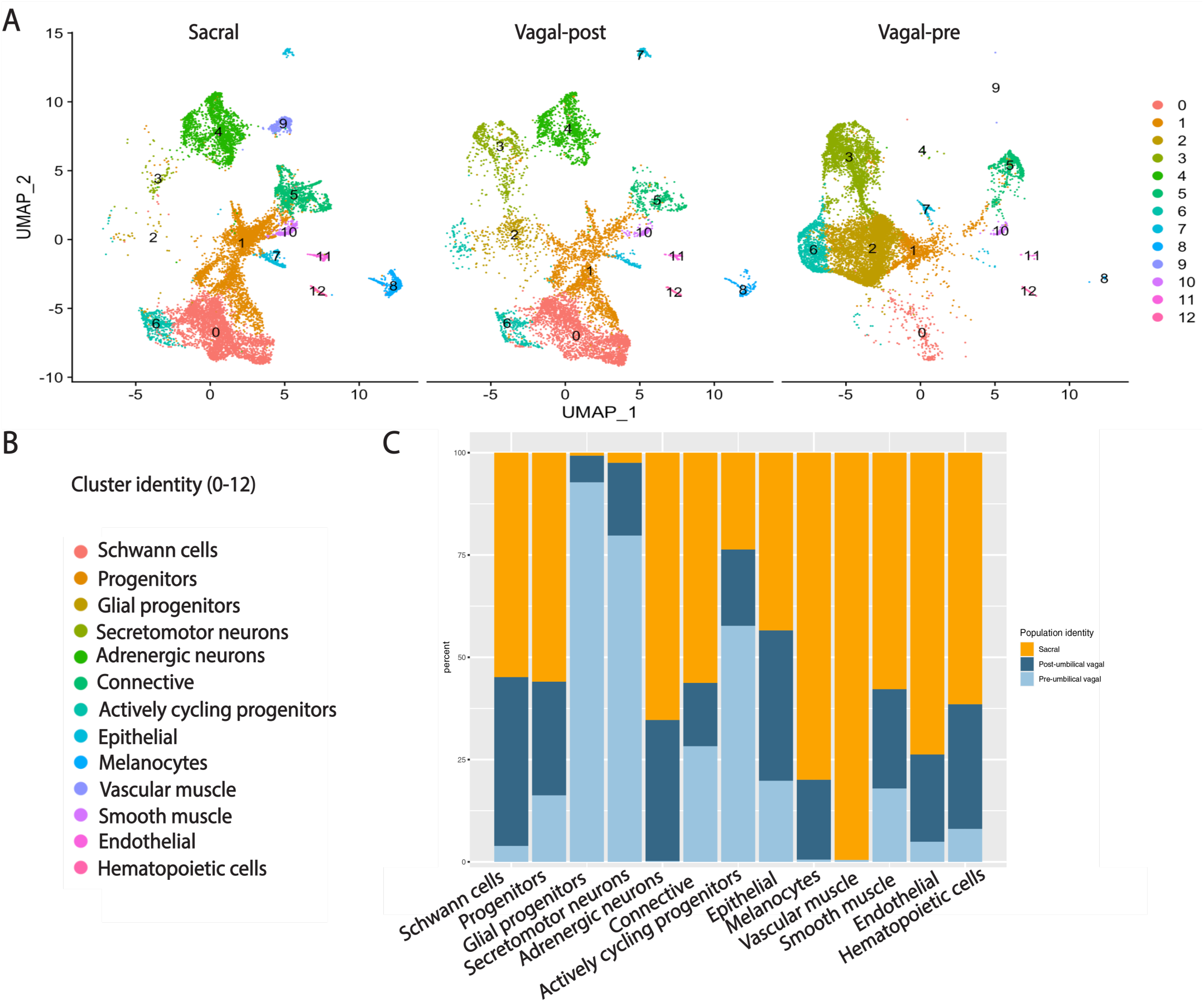
Sacral and vagal neural crest cells contribute to distinct ENS cell types. **(A)** UMAP separated by cell origin and position in the gut (sacral (red), sacral neural crest-derived cells in the post-umbilical region; vagal-post (green), vagal neural crest derived cells found in post-umbilical region; vagal-pre (blue), vagal neural crest derived cell located in the pre-umbilical region). **(B)** Cluster identities shown in (A). **(C)** Proportions of sacral, vagal-post, vagal-pre contribution across the clusters.

In summary, the main cell types derived from the sacral neural crest cells are Schwann cells, including those in intestinal nerve of Remak, cholinergic, adrenergic/serotonergic neurons, and melanocytes. Vagal neural crest-derived cells in the pre-umbilical region form a combination of neuronal progenitor cells and secretomotor neurons. The high percent of progenitor cells contributed by vagal-pre population suggests that cell differentiation is in progress all along the gut even though enteric migration is complete. Vagal neural crest cells in the post-umbilical gut contribute to most cell fates found in the ENS and form intermediate levels of cells between the pre-umbilical and sacral (Fig 4A, B). These results indicate that for those vagal neural crest cells that migrate long distances, environmental cues may have a profound influence on cell fate choice.

### Validation of marker expression by vagal versus sacral neural crest using dual retroviral lineage tracing

We next sought to validate the gene expression differences identified by single cell RNA-seq between sacral and vagal neural crest contributions to the gut in differentially labeled cell populations. To this end, we utilized axial-level specific retroviral labeling to sequentially mark vagal or sacral neural crest cells with different fluorophores in the same embryo. For identifying the vagal neural crest, RIA retrovirus expressing nuclear H2B-RFP was injected into the neural tube adjacent to somite 1-7. Embryos were then allowed to develop until HH17, at which time RIA retrovirus carrying H2B-YFP was injected into the neural tube posterior to somite 28 to label the sacral neural crest. The entire length of the gut was dissected and removed for immunohistochemistry at E10 and stained with antibodies to gene products identified as differentially expressed in our sc RNA-seq dataset (Fig 1A). To gain insight into the relative contributions of vagal versus sacral neural crest to the ENS along the entire length of the gut, we imaged transverse sections of six subregions along the anterior-posterior axis: esophagus (eso), stomach (sto), pre-umbilical small intestine (pre. int), post-umbilical small intestine (post.int), cecum (ce) and colon (col) (Fig 1A).

The results show that the pre-umbilical gut contained only H2B-RFP+, but no H2B-YFP+, labeled cells, suggesting that only vagal neural crest cells contributed to the pre-umbilical region (Fig 1A). This is consistent with previous studies using quail-chick chimeric systems^4^. Immunohistochemistry revealed vagal RIA retrovirus-labeled cells that co-expressed acetylcholine receptor (Fig 5A-C) and HuC/D (ELAV) in the pre-umbilical (Fig 5D-F), consistent with differentiated neurons. H2B-RFP and P0 double-positive Schwann cells were present along the pre-umbilical region (Fig 5G-I), as well as enteric progenitors or glial cells as determined by Sox10 expression (Fig 5J-L). Consistent with the scRNA-seq, there were only a few sparsely distributed neurons marked by TH (Fig 5M-O) and DBH (Fig 5P-R), TH+ cells were not observed in the pre-umbilical small intestine (Fig 5O) and DBH+ cells were absent from the esophagus (Fig 2P).

**Figure 5.**
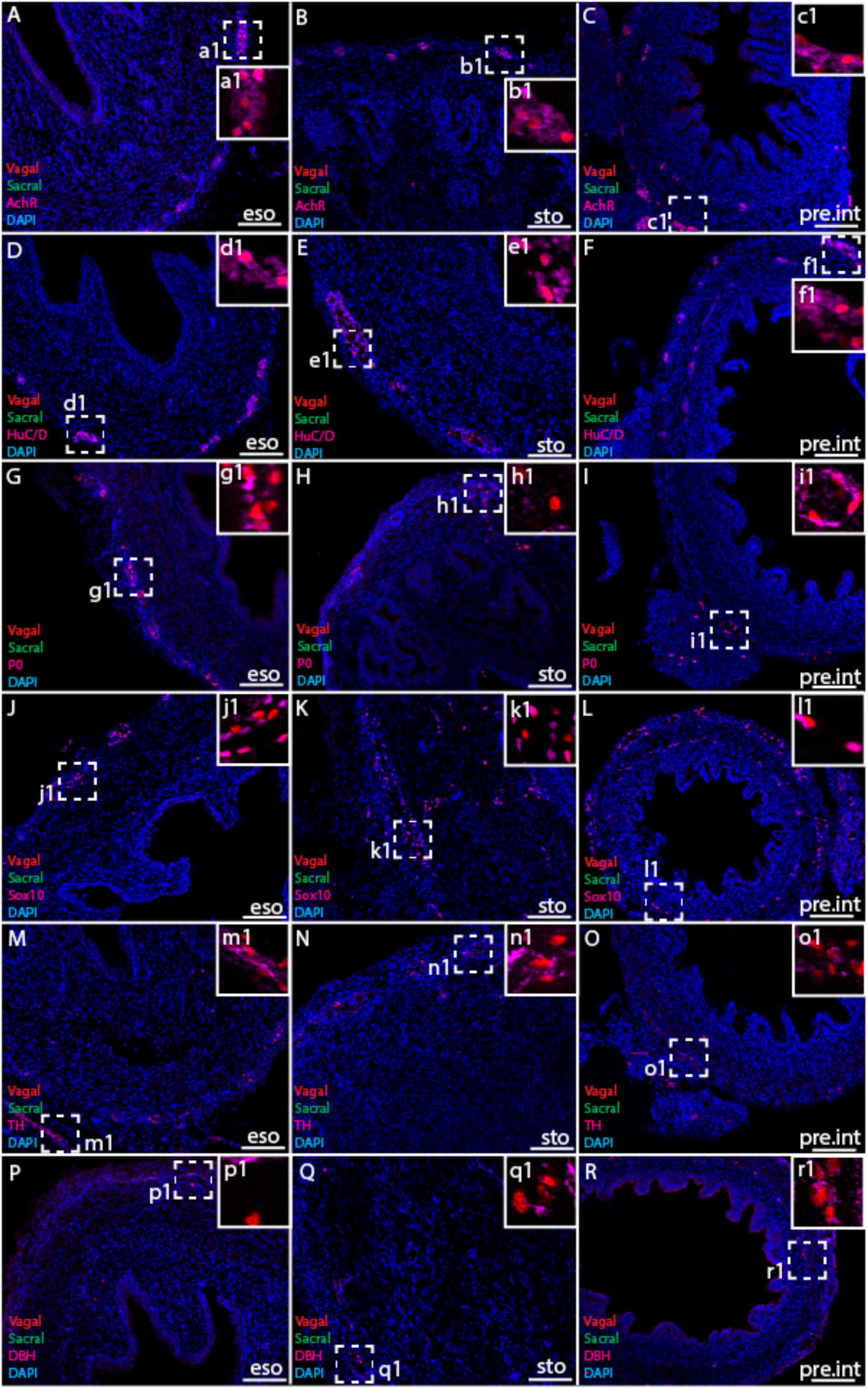
Vagal but not sacral neural crest contributes to pre-umbilical gut. **(A-C)** Acetylcholine receptor was expressed in some vagal neural crest-derived cells along the esophagus (eso, A), stomach (sto, B) and pre-umbilical small intestine (pre.int, C). **(D-F)** Vagal neural crest also gave rise to HuC/D + neurons in these regions. **(G-I)** Schwann cells were present in greater numbers in the esophagus (G) and pre-umbilical intestine (I) than in the stomach (H). **(J-L)** Enteric progenitor or glial cells expressing Sox10 in the nucleus were present in all three subregions. **(M-R)** A small number of neurons expressing TH (M-O) and DBH (P-R) were observed sparsely distributed in the pre-umbilical region. TH+ cells were not observed in the small intestine (O). DBH was absent in more anterior regions such as the esophagus (P) and stomach (Q). Insets a1-r1 show magnified regions in the corresponding dashed box. Sacral neural crest cells were absent from the pre-umbilical gut. Scale bars 160μm.

In contrast to the pre-umbilical gut, the post-umbilical ENS contained both H2B-RFP+ cells and H2B-YFP+ cells, indicating a collective contribution from sacral neural crest and some post-umbilical vagal neural crest (Fig 1A). Consistent with our scRNA-seq data, neural crest cells from different axial origins appeared to contribute to distinct neuronal subtypes. Acetylcholine receptor (AchR)-positive cells were more abundant in the vagal-derived population throughout the post-umbilical region (Fig 6A-C, inset a1-c1) whereas AchR+ cells from sacral neural crest cells were only observed in the colon (Fig 6A-C, inset a2-c2). Both vagal (Fig 6D-F, inset d1-f1) and sacral (Fi6D-F, inset d2-f2) neural crest populations differentiated into HuC/D+ neurons, except in the cecum where HuC/D labeling was more restricted to vagally-derived cells (Fig 6F, inset f2). Both vagal and sacral neural crest contributed to Schwann cells and ENS progenitors/glia expressing Sox10 and P0 in all subregions of the gut (Fig 6G-I, P0; J-L, Sox10). However, TH+ cells were most frequently derived from sacral neural crest cells in the colon region (Fig 6M-O, inset n2). These results confirm that a very large proportion of the ENS in the colon is derived from the sacral neural crest. DBH+ cells were also more abundant in sacral neural crest-derived population (Fig 6P-R), especially in regions highly enriched in sacral neural crest cells, such as the colon (Fig 6Q, inset q2) and cecum (Fig 6R, inset r2). These results suggest that vagal and sacral neural crest cells in the post-umbilical gut are distinct rather than functionally redundant, particularly in the hindgut.

**Figure 6.**
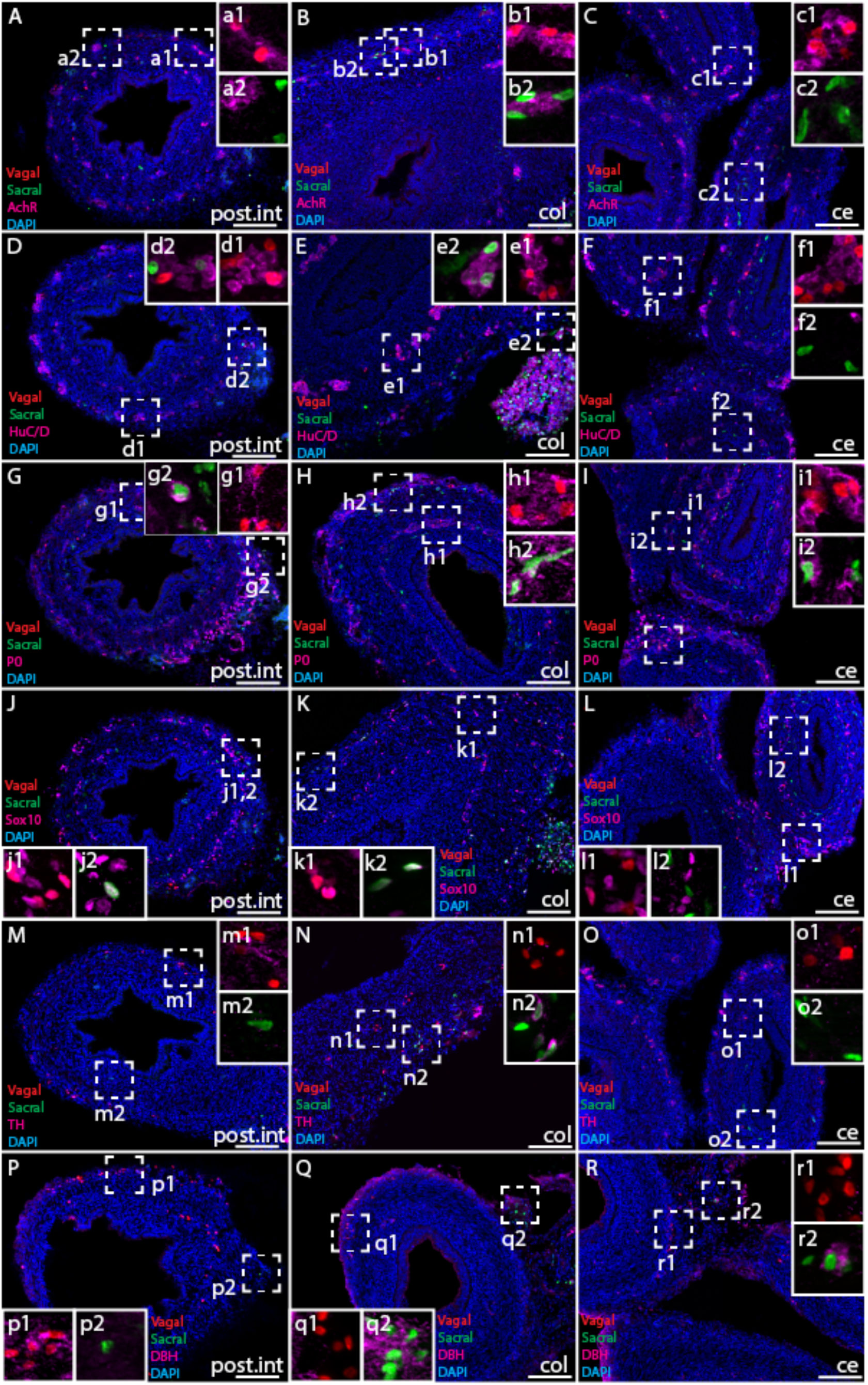
Relative contributions of vagal and sacral neural crest cells to the post-umbilical gut. **(A-C)** Acetylcholine receptor was broadly expressed by vagal neural crest along the post-umbilical gut (insets a1-c1); in contrast, sacral neural crest-derived cells did not express high level of AchR except in the colon region (insets a2-c2). **(D-F)** Differentiated neuronal marker HuC/D was expressed by vagal (insets d1-f1) and sacral (insets d2-f2) neural crest cells, except that sacral neural crest cells were absent from the cecum region (inset f2). **(G-L)** Schwann cells marked by P0, ENS progenitors/glia marked by Sox10 along the post-umbilical gut were derived from both populations. **(M-O)** TH+ cells were predominantly contributed by sacral neural crest cells in the colon region (inset n2). **(P-R)** Some vagal neural crest cells in the small intestine expressed DBH (inset p1), but most DBH+ cells were derived from sacral neural crest-derived in the colon (inset q2) and the cecum (inset r2). Insets a1-r1, a2-r2 show magnified regions in the corresponding dashed box. Scale bars 160μm.

### Sub-classification of vagal and sacral neural crest-derived neuronal cell types

To further probe for specific neurotransmitter characteristic within the chick ENS, we extracted all cells from secretomotor and neuronal clusters (Fig 7A’) and clustered the cells into 9 subsets (Fig 7A, inset of Fig 7A’). We examined a variety of receptors, neurotransmitters and neuropeptides that mark specific neuronal cell fates^19^ and plotted the expression in the subsets (Fig 7B).

**Figure 7.**
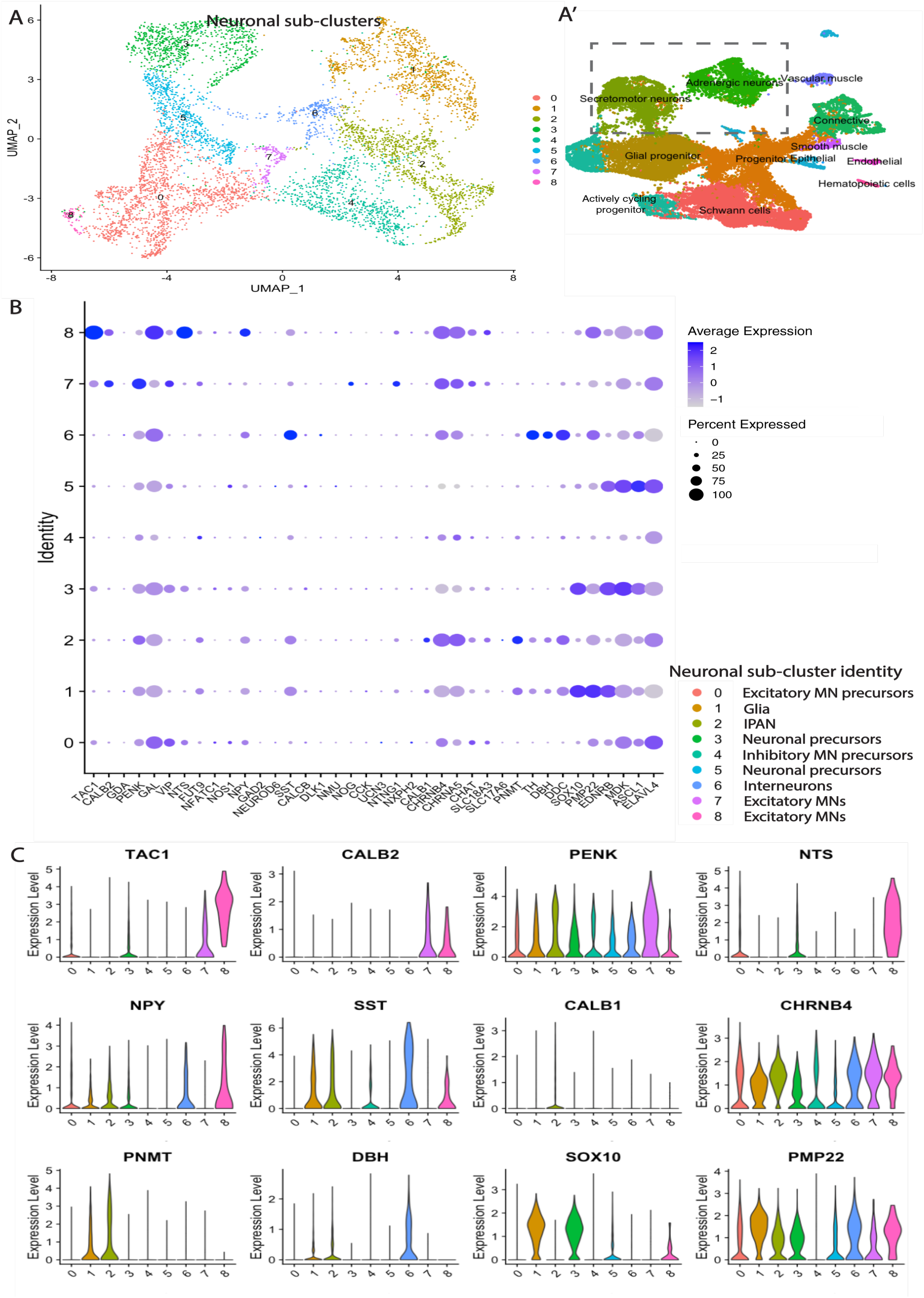
Sub-classification of neuronal clusters. **(A)** Sub-clustering of neuronal clusters (secretomotor, neuronal) found in A’ resulted in 9 distinct populations. **(B)** Dot plot representing neurotransmitters, neuropeptides, receptors and key genes representing specific neuronal/glial functions, classifying the clusters into excitatory motor neurons, glia, IPANs, inhibitory motor neurons, interneurons and their precursors not yet fully differentiated. **(C)** Violin plot representing neurotransmitters, neuropeptides, receptors and key genes representing specific neuronal/glial functions. IPANs: intrinsic primary afferent neurons.

C1 of this subsetting is defined as a glial cluster due to its high and broad expression of *Sox10*, *Pmp22* and *Ednrb* (Fig 7B, Fig 7C *Sox10*). C3 shares some similarities with C1, most likely reflecting neuronal precursors as this cluster has lower *Pmp22* and higher *Mdk* expression (Fig 7B). C0 appears to be a precursor for excitatory motor neurons, due to its relatively high expression of *Tac1*, *Penk*, *Gal* and *Vip*, with some cholinergic transcripts and moderately high level of *Mdk* (Fig 7B, Fig 7C *Tac1*, *Penk*). In contrast, C7 is identified as excitatory motor neuron due to abundant expression of *Calb2*, *Penk* and *Vip* with possibly a small number of intrinsic primary afferent neurons (IPANs), identified by *Nog* and *Ntg1* expression (Fig 7B, Fig 7C CALB2, PENK). C8, a *Tac1*, *Gal*, *Nts* positive cluster, is also identified as excitatory motor neurons (Fig 7B, Fig 7C *Tac1*, *Nts*). C2 is likely to be a combination of excitatory motor neurons and CALB1+ IPANs, with cholinergic/catecholaminergic properties (Fig 7B, Fig 7C *Calb1*).

C4 lacks a definitive cell marker, but a small percentage of cells show high expression of the inhibitory motor neuron marker *Fut9* (Fig 7B). Other inhibitory motor neuron markers remain low, suggesting these cells may not be fully differentiated due to low *Elavl4* expression (Fig 7B). Accordingly, we ascribed it as inhibitory motor neuron precursor. C5 is another cluster of differentiating neurons based on *Ascl1* expression. C6, in contrast, is the cluster with most *Sst* expression, indicating an interneuron cell fate (Fig 7B, Fig 7C *Sst*). C6 is also likely to be the major sub-cluster with catecholaminergic properties found specifically in post-umbilical gut (Fig 7C *Pnmt*, *Dbh*).

Together, these results confirm the differential gene analysis in post-umbilical vagal and sacral bulk transcriptome analysis. In general, vagal neural crest cells are the primary source of motor neurons, as indicated by upregulation of *Calb2* and *Nts* in vagal-post, whereas sacral neural crest cells give rise to adrenergic and serotonergic neuronal types, as reflected by high fold change in *Dbh*, *Th*, *Pnmt* and *Ddc* (Fig 1C, D).

## Discussion

The enteric nervous system regulates critical gastrointestinal functions including digestion, hormone secretion and immune interactions. Abnormal ENS development can lead to enteric neuropathies including Hirschsprung’s disease, characterized by lack of motility and obstruction. Studies have suggested a critical role for the vagal neural crest in the etiology of Hirschsprung’s disease^7, 11^. However, the role of the sacral neural crest, which colonizes the hindgut in close coordination with the vagal neural crest, has been largely understudied in enteric neuropathies. To better understand the derivatives of the sacral neural crest and their coordination with the vagal neural crest during ENS development, here we examine the diversity of cell types arising from vagal versus sacral axial levels at single cell resolution.

Ablation and heterotopic grafting experiments have previously been used to study the interplay between the vagal and sacral populations but have led to contradictory interpretations. While some studies concluded that vagal and sacral neural crest exhibit autonomous migration properties independent of the environment, others suggested a role for environmental influences. On the one hand, an aganglionic hindgut model created by surgically removing the caudal part of vagal neural crest and replacing it with quail sacral cells found that quail sacral cells migrated into the hindgut with a small increase in number of neurons. This suggested that sacral neural crest cells do not require the vagal population to migrate^8^. Reciprocally, when vagal neural crest cells are grafted to the sacral region, they migrate earlier and produce a larger neuronal population than the endogenous sacral neural crest cells^19^. However, other studies suggested a more prominent environmental effect, such that interchanged vagal and sacral neural crest cells migrated according to the local environment^20^. Consistent with this, combining chick gut before neural crest colonization with chick or quail neural crest revealed that sacral neural crest cells can colonize the colorectum independent of the vagal neural crest, but require the hindgut environment to differentiate^21^.

Our results using axial-level specific labeling provide a complementary approach to address these questions by comparing transcriptome between vagal and sacral neural crest for the first time. We show that sacral and vagal-derived ENS cell types are not equivalent. For example, we find *Tnc* expression to be specific to the vagal neural crest (Fig 1C, D), consistent with its requirement for migration into the hindgut by changing the extracellular microenvironment^22^. In addition, the sacral neural crest expresses high levels of *Sox10*-mediated *Cdh19*^23^, and well as *Pax3* and 5-HT3 for innervation and neuronal maturation in the pelvic ganglion^24, 25^. *Ret* is also known to be upregulated in vagal neural crest cells to mediate more invasive behavior than in sacral neural crest^26^. In the hindgut, distinct properties of vagal and sacral neural crest suggest that they are likely to play disparate roles in enteric neuropathies. Indeed, diverse timeframes and migration patterns expose the cells to differential environments, leading to divergent cell fates and functions.

Previous studies have provided valuable information regarding gene expression, cell fate divergence, gene regulatory networks and chromatin landscape of vagal-derived neural crest cells^14, 15, 27–29^. By focusing transcriptome analysis on the small intestine in mice at postnatal day 21, one study identified 12 distinct neuronal classes categorized by a combination of neurotransmitters; the authors found *Pbx3* to be a key gene for differentiation. Our results have identified most neurotransmitter genes found in the murine system with the exception of NOS1+ nitrergic neurons, GAD2+ GABAergic neurons, or SLC17A6+ Glutamatergic neurons^14^, which may develop at a later time point than studied here (Fig 7B). Another study, utilizing RAISIN RNA-seq, identified 21 neurons and 3 glial clusters in the mouse small intestine and colon according to subsets of neurotransmitters. Neuronal and glial subsets we identified generally agree with this, showing an overlapping group of sensory neuron markers such CCK, VIP, SST, NOG and NMU^29^ (Fig 7B). Compared with these studies done with postnatal and adult tissue, we observed more clusters with progenitor/precursor identity (Fig 2C, Fig 7B), which is not unexpected, given that our analysis utilizes the gut from embryonic stages. While previous studies did not separate vagal from sacral contributions, the small intestine and colon are likely to be composed of both vagal-post and sacral populations, which are distinct from anterior vagal derivatives. Our analysis reveals that gene expression patterns are markedly different according to axial-level, confirming the results from previous studies within proximal versus distal colon^29^. This important conclusion suggests that the ENS is not uniformly distributed throughout the gut but varies from proximal to distal.

Taken together, the present results suggest that there are different developmental programs for vagal versus sacral neural crest population. Our results help explain sacral neural crest cannot completely compensate for the loss of vagal neural crest. The cell composition of the post-umbilical ENS is distinct from that of pre-umbilical ENS, with major differences largely resulting from the differential contributions of the sacral neural crest. In addition, the differentiation program of vagal-derived neural crest in the post-umbilical gut is different from that of the pre-umbilical vagal population, suggesting that environmental factors have a large influence on cell fate. Our study highlights the contributions of the sacral neural crest, particularly to the hindgut, the region most affected in Hirschsprung’s disease characterized by colonic agangliogenesis.

## Methods

### Retroviral labeling and chick embryology

H2B-YFP (#96893) and H2B-RFP (#92398) obtained from Addgene were cloned into the RIA plasmid between Not1 and Cla1 sites. RIA-H2B-YFP/RFP was transfected into DF1 cells (ATCC, Manassas, VA; #CRL-12203, Lot number 62712171, Certificate of Analysis with negative mycoplasma testing available at ATCC website) using PEI standard transfection protocol. DF1 cells were maintained in Gibco Dulbecco’s Modified Eagle Medium (DMEM) supplied with 10% FBS for 4 days, with 12ml of supernatant collected per day. The supernatant was concentrated using ultracentrifuge for at 76000g for 1.5 hours to get a viral stock tittered about 10^7^ pfu/ml, aliquoted and stored at −80 °C until use. Viral solution was supplemented with 0.3μl of 2% food dye (Spectral Colors, Food Blue 002, C.A.S# 3844-45-9) as an indicator, injected to fill the neural tube between somite 1-7 at HH Stage 10 to label vagal neural crest and/or posterior to somite 28 at HH Stage 17 to label sacral neural crest *in ovo*. Embryos were supplied with Ringer’s Solution (0.9% NaCl, 0.042%KCl, 0.016%CaCl_2_ • 2H_2_O wt/vol, pH7.0), sealed with surgical tape, and incubated at 37 °C until embryonic day 10.

### Gut cell dissociation and Fluorescence-Activated Cell Sorting (FACS)

At embryonic day 10, the gastrointestinal tract was dissected from chick embryos and washed with Ringer’s solution. Pre- and post-umbilical regions were separated, broke into pieces in chilled DPBS and loose-fit homogenized in Accumax solution (EMD Millipore). 400μl of Accumax-tissue mixture was aliquoted into 1.7 ml Eppendorf tubes and shaken at 37 °C for 12mins. After dissociation, chilled Hanks Buffered Saline Solution (HBSS) supplemented by BSA (125mg in 50ml, Sigma; 0.2% w/v) and 1M HEPES (500μl in 50ml, PH7.5, ThermoFisher) was added to quench the reaction. The dissociated cells were passed through a 70μm cell strainer (Corning) and collected by centrifuging at 500g for 11mins at 4 °C. The cells were resuspended in HBSS-BSA, supplemented 7-AAD Viability Staining Solution (Biolegend # 420404, 500 TESTS), and sorted for YFP+, viable single cells using Sony SY3200 cell sorter at the Caltech Flow Cytometry Facility.

### Bulk transcriptome analysis

For vagal neural crest in the post-umbilical regions, sacral neural crest at post-umbilical regions, 2 biological replicates were processed, with each replicate containing YFP+ cells from 3 embryos. 2000 cells per replicate were lysed to generate cDNA library using SMART-Seq v4 Ultra Low Input RNA Kit (Takara Bio). The library was sequenced with 50 million single end reads with 50 bp length using HiSeq 2500 at the Millard and Muriel Jacobs Genetics and Genomics Laboratory Caltech. Sequencing reads were trimmed using *cutadapt*^30^ and mapped to Galgal6 genome using *Bowtie2*^31^. *DESeq2*^32^ analysis was performed to find differential expressed genes between vagal and sacral neural crest at post-umbilical regions generated by *HTseq-count*^33^. Differential gene expression was presented using Volcano Plot (coloring genes with Fold Change>2 and p value<0.05) and Heatmap2 provided by the Galaxy platform. Because there were more upregulated genes in the sacral than vagal-post populations, we annotated genes related to neuronal function for sacral population, genes related to neuronal function as well as genes with top fold-change and top significance in vagal-post population.

### Single-cell transcriptome analysis and data processing

For vagal neural crest in pre- and post-umbilical gut, sacral neural crest in post-umbilical gut, 2 biological replicates were obtained. Each replicate of pre-umbilical gut was pulled from 3 embryos; each replicate containing post-umbilical gut was pulled from 6 embryos. After FACS for viable YFP+ cells, 4600-5000 cells per replicate were used for library preparation by the SPEC at Caltech. The library was sequenced on NovaSeq S4 lane with 2×150bp reads by Fulgent Therapeutics. To process fastq raw data, standard ENSEMBL galgal6 reference database was used. Single-cell level gene quantification was then performed using *Cellranger* v3.1.0^16^ and *kallisto* 0.46.2 & *bustools* 0.40.0 pipelines^17^ with default parameters. Gene count matrices from all the samples were combined and only cells with more than 200 genes detected were kept for the downstream analysis. To further remove potential doublet cells, *DoubletFinder* 2.0.3 package was used. Gene counts were normalized and scaled using Seurat v3.2.0^18^. The first 30 principal components from PCA analysis were used to find neighbors with *Findneighbors* function before cell clustering with *FindClusters* function (resolution = 0.2). UMAP dimensionality reduction was performed using RunUMAP function with uwot-learn selected for the parameter umap.method.

### Immunohistochemistry and imaging

Gastrointestinal tracts were dissected and fixed in 4% PFA in PBS (PH7.5) for 25 mins at 4°C and washed with PBS for three times. Pre- and post-umbilical regions were separately incubated in 15% sucrose at 4°C overnight and in gelatin at 37°C for 2 hours. Gut segments were embedded in gelatin solution, flash-frozen with liquid nitrogen, and mounted with Tissue-Tek O.C.T compound (Sakura #4583) for sectioning (Microm HM550 cryostat). Gut sections were incubated in 1xPBS at 42°C until the gelatin was dissolved, soaked in 0.3% vol/vol Triton-X100 in 1xPBS for permeabilization. Blocking buffer was prepared in 1xPBS with 5% vol/vol normal donkey serum and 0.3% vol/vol Triton-X100. Sections were incubated with primary antibody at 4°C overnight Sections were washed with 1xPBS for 10 minutes and 3 times. After the washed, sections were incubated with secondary antibody for 45 minutes at room temperature. List of primary antibodies used: 1:20 chicken anti AchR ratIgG2a, mAB270 (DSHB Antibody Registry ID: AB_531809); 1:500 Mouse anti HuC/D IgG2b (Invitrogen-Cat#A21271); 1:20 Chicken anti mouse P0 IgG1, IE8 (DSHB Antibody Registry ID: AB_2078498); 1:500 Rabbit anti Sox10 (Millipore-Cat# AB5727); 1:500 Rabbit anti Tyrosine Hydroxylase (Millipore-Cat# AB152); 1:500 Rabbit anti DBH (Immuno Star-Cat #:22806). List of secondary antibodies used: 1:1000 donkey anti-mouse IgG2b 647 (Invitrogen A31571), 1:1000 goat anti-mouse IgG1 647 (Invitrogen A21240), 1:1000 donkey anti-rat IgG 647 (Abcam ab150155), 1:1000 goat anti-rabbit IgG 647 (Invitrogen A21245). Sections were imaged with Zeiss AxioImager.M2 with Apotome.2. Images were cropped and magnified for representation.

## Acknowledgement

This work was supported by R01DE027568 and R35NS111564 to M.E.B. We thank Drs. Igor Antoshechkin and Vijaya Kumar and the Millard and Muriel Jacobs Genetics and Genomics Laboratory at California Institute of Technology for their guidance and support in bulk RNA-sequencing. We thank Jamie Tijerina and Rochelle Diamond from the Beckman Institute Flow Cytometry Facility for their help with the FACS. We thank Dr. Sisi Chen, Jeff Park, Prof. Matt Thomson and SPEC at Caltech for their dedicated support in optimization and guidance in single-cell RNA-sequencing. We thank Dr. Fan Gao and Bioinformatics Resource Center in the Beckman Institute at Caltech for guiding us through single-cell transcriptomic analysis. We appreciate the help from Prof. Carlos Lois for kindly sharing equipment with us to perform RIA concentration. We thank Dr. Michael L. Piacentino, Dr. Erica J. Hutchins and Prof. Angelike Stathopoulos for the helpful discussion on the manuscript.

